# Production of functional oocytes requires maternally expressed *PIWI* genes and piRNAs in golden hamsters

**DOI:** 10.1101/2021.01.27.428354

**Authors:** Hidetoshi Hasuwa, Yuka W. Iwasaki, Au Yeung Wan Kin, Kyoko Ishino, Harumi Masuda, Hiroyuki Sasaki, Haruhiko Siomi

**Affiliations:** Department of Molecular Biology, Keio University School of Medicine, Tokyo, Japan; Division of Epigenomics and Development, Medical Institute of Bioregulation, Kyushu University, Fukuoka, Japan; Japan Science and Technology Agency (JST), Precursory Research for Embryonic Science and Technology (PRESTO), Saitama, Japan

## Abstract

Many animals have a conserved adaptive genome defense system known as the Piwi-interacting RNA (piRNA) pathway which is essential for germ cell development and function. Disruption of individual mouse *Piwi* genes results in male but not female sterility, leading to the assumption that *PIWI* genes play little or no role in mammalian oocytes. Here, we report generation of *PIWI*-defective golden hamsters, which reveals defects in the production of functional oocytes. The mechanisms involved vary among the hamster *PIWI* genes; lack of *PIWIL1* has a major impact on gene expression, including hamster-specific young transposon de-silencing, whereas *PIWIL3* deficiency has little impact on gene expression in oocytes, although DNA methylation was found to be reduced to some extent in *PIWIL3*-defecient oocytes. Our findings serve as the foundation for developing useful models to study the piRNA pathway in mammalian oocytes, including humans, which is not possible with mice.

## Introduction

piRNAs are derived from specific genomic loci termed piRNA clusters and form effector complexes with PIWI proteins, a germline-specific class of Argonaute proteins, to guide recognition and silencing of their targets, mostly transposable elements (TEs) (1–3). Mammalian PIWI-piRNA pathways have mostly been studied in mice, which express three PIWI proteins (MIWI/PIWIL1, MILI/PIWIL2, and MIWI2/PIWIL4) abundantly in the testis, but only weakly in the ovary (1–3). These PIWI proteins bind distinct classes of piRNAs which direct chromatin modifications of target TE sequences during embryogenesis and guide silencing of target TEs at the posttranscriptional level later in spermatogenesis, to ensure completion of meiosis and successful sperm production. Deficiency of *Piwi* genes in mice is characterized by spermatogenesis arrest and infertility in males, but females with deficiencies in these genes remain fertile (4–9) with limited impact on TE silencing (10, 11). These findings clearly demonstrate that mouse *Piwi* genes are essential for male, but not female, germ cell development.

However, most mammalian species including humans possess an additional *PIWI* gene, termed *PIWIL3*, in addition to the three *PIWI* genes described above (12, 13). Thus, piRNA-mediated silencing may differ between mice and other mammals with four *PIWI* genes. However, little is known about the piRNA pathway in mammalian species with the four *PIWI* genes, particularly their potential roles in functional oocyte production. Golden Syrian hamsters (golden hamster, *Mesocricetus auratus*) have been used as an experimental rodent model for studying human diseases, particularly, cancer and infectious diseases, including the recent Covid-19, because they display physiological and pharmacological responses resembling those of humans (14, 15). In addition, genome editing using the CRISPR/Cas9 system was recently enabled in golden hamsters to engineer the genes of interest (16, 17). Unlike laboratory mice and rats, which belong to the *Muridae* family of rodents lacking *PIWIL3*, the golden hamster belongs to the *Cricetidae* family of rodents and has four distinct *PIWI* genes. We recently found that hamster *PIWIL1* and *PIWIL2* are expressed in both the testis and the ovary, whereas *PIWIL3* and *PIWIL4* are exclusively expressed in the ovary and testis, respectively (18).

## Result

In the present study, we aimed to investigate the roles of *PIWI* genes in female reproduction by studying *PIWIL1* and *PIWIL3* genes, both of which were highly expressed in oocytes (18; see also Fig. S3A and B). We employed direct injection of Cas9 mRNA or Cas9 protein together with sgRNAs into the pronucleus (PN) and/or cytoplasm of an embryo to generate *PIWIL1*- and *PIWIL3*-deficient hamsters. We injected the cells under a microscope with red filters (600 nm) in a dark room, because even brief exposure of hamster zygotes to light results in total developmental arrest (16). Injected embryos with a normal morphology were transferred to each oviduct (10–15 embryos per oviduct) of pseudo-pregnant recipient females. In total, 46 pups were obtained from the *PIWIL1*-sgRNA injected embryos, of which 23 carried a mutant allele, as demonstrated by genomic PCR and sequencing using the tail tissue (Fig. S1). However, nine female founder hamsters were infertile and could only establish three frameshift mutant hamster lines. For the production of *PIWIL3*-deficient hamsters, 18 pups were obtained from the injected embryos. Genomic sequencing revealed some mosaicism and identified a total of 13 types of *PIWIL3* mutant alleles (Fig. S2). We crossed F1 heterozygous mutant hamsters to generate F2 homozygous mutant hamsters, which yielded offspring with genotypes that segregated in a Mendelian distribution. Western blots confirmed the lack of PIWIL1 or PIWIL3 in the mutant ovaries. Immunostaining also demonstrated the loss of PIWIL1 or PIWIL3 protein in the cytoplasm of mutant growing oocytes (Fig. S3A and B).

Both *PIWIL1*- and *PIWIL3*-deficient hamsters appeared normal without outwardly discernible morphological and behavioral abnormalities. *PIWIL1*-deficient male hamsters displayed small testes and lack of mature sperm in the cauda epididymis (Fig. S4A-E). DDX4 (mouse Vasa) and acrosome staining with lectin peanut agglutinin (PNA) revealed spermatogenesis arrest in the pachytene stage and a complete lack of S2 spermatids in the *PIWIL1* mutant testis (Fig. S4F and G). These developmental abnormalities resembled those observed in *Miwi*/*Piwil1*-deficient male mice (4). *PIWIL3* was not expressed in the testis and *PIWIL3*-deficient male hamsters displayed no overt phenotype. Histological examination of ovaries from *PIWIL1*- and *PIWIL3*-deficient adult females revealed no gross abnormalities (Fig. S3C). *PIWIL1*- and *PIWIL3*-deficient female hamsters displayed complete sterility and reduced fertility, respectively (Fig. 1). When *PIWIL1*-deficient females were crossed with heterozygous males, they never became pregnant despite successful coitus demonstrated by the presence of sperm in the vagina (Fig. 1A and B). *PIWIL3*-deficent females, when mated with heterozygous males, displayed reduced fertility with both reduced pregnancy rate (43.5% versus 75% in homozygous and heterozygous females, respectively) and smaller litter size (4.6 versus 6.8 in homozygous and heterozygous females, respectively) (Fig. 1A and B). The homozygous *PIWIL3* mutant pups proceeded into adulthood without expressing any overt developmental abnormalities.

**Fig. 1.**
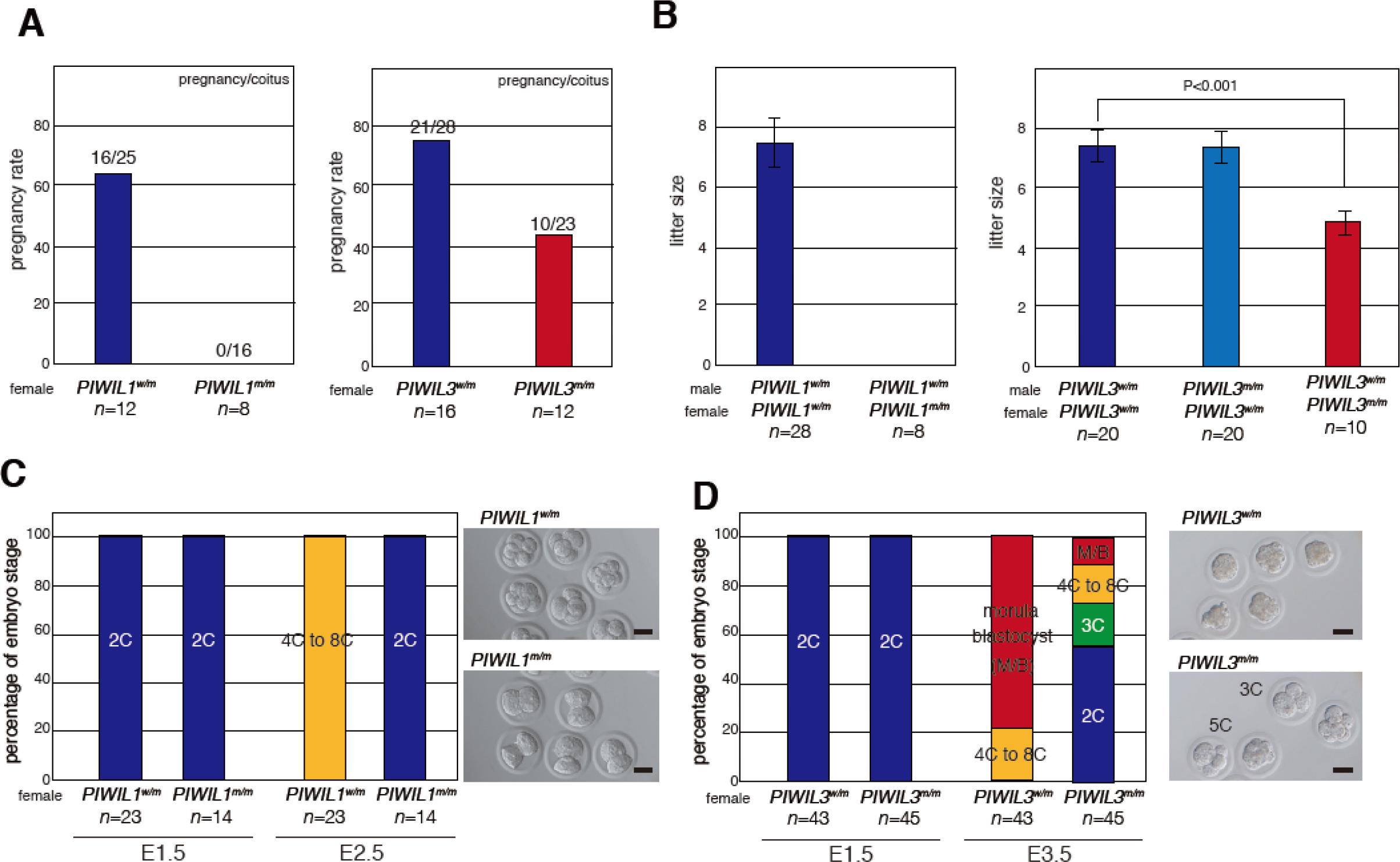
Female fertility phenotypes of *PIWIL1* and *PIWIL3* mutant golden hamsters. **(A)** Fecundity of female golden hamsters was analyzed by natural mating; *n* indicates the number of female hamsters used in each genotype. The numbers on the bar indicate pregnancy per coitus. w/m: heterozygous, m/m: homozygous mutants. **(B)** Litter sizes from mated females; *n* shows the number of litters analyzed, and error bars indicate ± s.e.m. **(C, D)** The early embryogenesis ratio was analyzed *in vitro*. 2C embryos from *PIWIL1* heterozygous and homozygous mutant female hamsters mated with wild-type males were cultured for 1 d *in vitro* **(C)**. 2C embryos from *PIWIL3* heterozygous and homozygous mutant female hamsters mated with wild-type male were cultured for 2 d *in vitro* **(D)**. Images of representative embryos (at E2.5 in **C** and at E3.5 in **D**) with the indicated genotypes are shown on the right. Note three-cell or five-cell embryos in the culture of *PIWIL3* mutant embryos.

To examine how embryogenesis proceeds after fertilization with *PIWIL1*- or *PIWIL3*-deficient oocytes, we isolated two-cell (2C) embryos from *PIWIL1*- or *PIWIL3*-deficient females crossed with wild-type males and cultured them *in vitro*. All maternal *PIWIL1*-deficient 2C embryos remained at the 2C stage and apparently died after one day of in vitro culture, while 2C embryos isolated from heterozygous females developed into four-cell to eight-cell embryos under the same culture conditions (Fig. 1C). A high proportion (55%) of maternal *PIWIL3*-deficient 2C embryos showed a 2C arrest phenotype even after two days of in vitro culture and a significant portion of the others were arrested at the three- or five-cell stages, indicating that one blastomere of 2C was arrested and the other divided (Fig. 1D). These results indicate that both *PIWIL1*- and *PIWIL3*-deficent oocytes can be fertilized and can proceed through the first cell division, but they fail to develop, either completely or partially, at subsequent developmental stages. These results also highlight the non-redundant essential role of *PIWI* genes in hamster oogenesis.

To gain insight into the molecular defects leading to the observed abnormalities, we performed small RNA sequencing (small RNA-seq) of *PIWIL1*- and *PIWIL3*-deficient oocytes; results revealed a decrease in specific populations of small RNAs (Fig. 2A). In both heterozygous control oocytes, three peaks at 19 nucleotide (nt), 23 nt, and 29 nt were observed. We compared the small RNA length distribution between heterozygous and homozygous oocytes by normalizing the whole small RNA population using the detected miRNA reads. This revealed a significant decrease in the population of 23- and 29-nt small RNAs in *PIWIL1*-deficient oocytes. In contrast, only the 19-nt small RNA population was decreased in *PIWIL3*-deficient oocytes. This is consistent with our recent findings that PIWIL1 binds to two populations of small RNAs, 23 and 29 nt (PIWIL1-bound piRNAs), and PIWIL3 binds to 19-nt small RNAs (PIWIL3-bound piRNAs) in metaphase II (MII) oocytes (18). It has been recently shown that human PIWIL3 binds a class of ∼20-nt small RNAs in oocytes (19), suggesting that hamster PIWIL3 resembles the human ortholog.

**Fig. 2.**
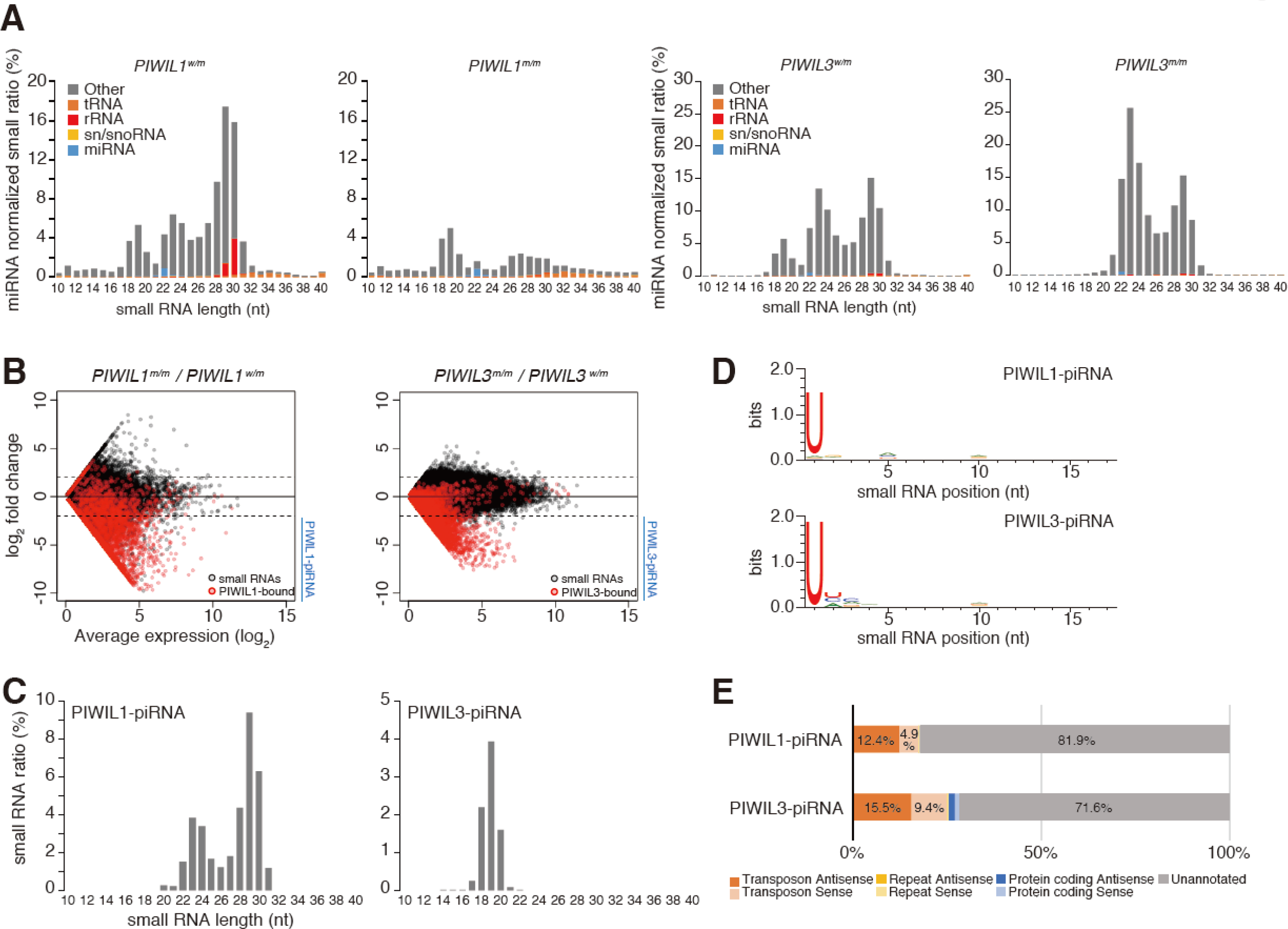
Characterization of small RNAs in oocytes of *PIWIL1*- or *PIWIL3*-deficient hamsters. **(A)** Composition of small RNA categories according to their length distributions in hamster MII oocytes from *PIWIL1* mutants (left) and *PIWIL3* mutants (right). Small RNA reads from known ncRNAs (tRNA, rRNA, sn/snoRNA, miRNA) are indicated by the described colors. The reads were normalized to the reads mapped to miRNAs, and the total number of obtained reads. **(B)** An MA plot showing the expression level of each unique small RNA sequence. To calculate log2 fold change and expression levels, 1 was added to each value to calculate the levels of small RNA sequences with 0 in either of the samples. Small RNAs identical to the PIWIL1-bound piRNAs/PIWIL3-bound piRNAs (16) and those that decreased by >4-fold in homozygous mutants were defined as PIWIL1-piRNA or PIWIL3-piRNA (indicated by blue lines). The red dots indicate piRNA bound to PIWIs described previously (18). **(C)** The size distribution of PIWIL1-piRNA and PIWIL3-piRNA. Obtained small RNA reads were normalized to the total number of genome-mapped small RNA reads. **(D)** WebLogo analysis of PIWIL1-piRNA and PIWIL3-piRNA. **(E)** Annotation of genome mapped PIWIL1-piRNA and PIWIL3-piRNA.

To further characterize the decreased population of small RNAs, changes in the expression level of each small RNA were analyzed in *PIWIL1* and *PIWIL3* mutants. A decrease in expression level was observed in populations of small RNAs (Fig. S5), consistent with our observation of the length distribution (Fig. 2A). Comparisons with the previously identified PIWIL1-bound/PIWIL3-bound piRNAs (18) indicated that the small RNAs, the populations of which were found to decrease, were identical to these PIWIL1/PIWIL3-bound piRNAs (Fig. 2B), showing that the lack of PIWI proteins depleted piRNAs that bind to them. In contrast, the expression levels of a minor population of the PIWIL1/PIWIL3-piRNAs were not decreased in homozygous mutants. This may be because these piRNAs can be bound by other PIWI proteins and/or can be stably present in a PIWI-unassociated manner. Small RNAs identical to the PIWIL1-bound piRNAs/PIWIL3-bound piRNAs (18) and those that decreased by over 4-fold in homozygous mutants were defined as PIWIL1-piRNA or PIWIL3-piRNA. As expected, PIWIL1-piRNAs were mostly 23- and 29-nt small RNAs, and PIWIL3-piRNAs were mostly 19-nt small RNAs (Fig. 2C). They both possessed uracil (U) at their 5’ end, which is a conserved characteristic of piRNAs (1, 2) (Fig. 2D). Genome mapping and annotation of PIWIL1- and PIWIL3-piRNAs revealed that they were mainly mapped to unannotated regions of the genome, as in the case of mouse pachytene piRNAs (3). A total of 12.4% PIWIL1-piRNAs and 15.5% of PIWIL3-piRNAs were mapped to the antisense orientation of TEs, suggesting that they could target TEs (Fig. 2E).

We also performed RNA sequencing (RNA-seq) using samples isolated from *PIWIL1*- and *PIWIL3*-deficient MII oocytes. This revealed that the lack of *PIWIL1* had a significant impact on the oocyte transcriptome (Fig. 3A). We detected 1612 differentially expressed genes (DEGs) in *PIWIL1*-deficient oocytes, 66.13% of which were up-regulated, indicating possible silencing of these genes by the PIWI-piRNA pathway. However, only 0.02% of the PIWIL1-piRNAs were mapped to the antisense direction of protein-coding genes (Fig. 2E), suggesting that they cannot directly target the mRNA of these DEGs. In contrast, the level of most genes remained unchanged in *PIWIL3*-deficient oocytes, with only 21 DEGs detected (Fig. 3A). These results indicate that a number of genes are regulated by *PIWIL1*, but not *PIWIL3*, in hamster oocytes.

**Fig. 3.**
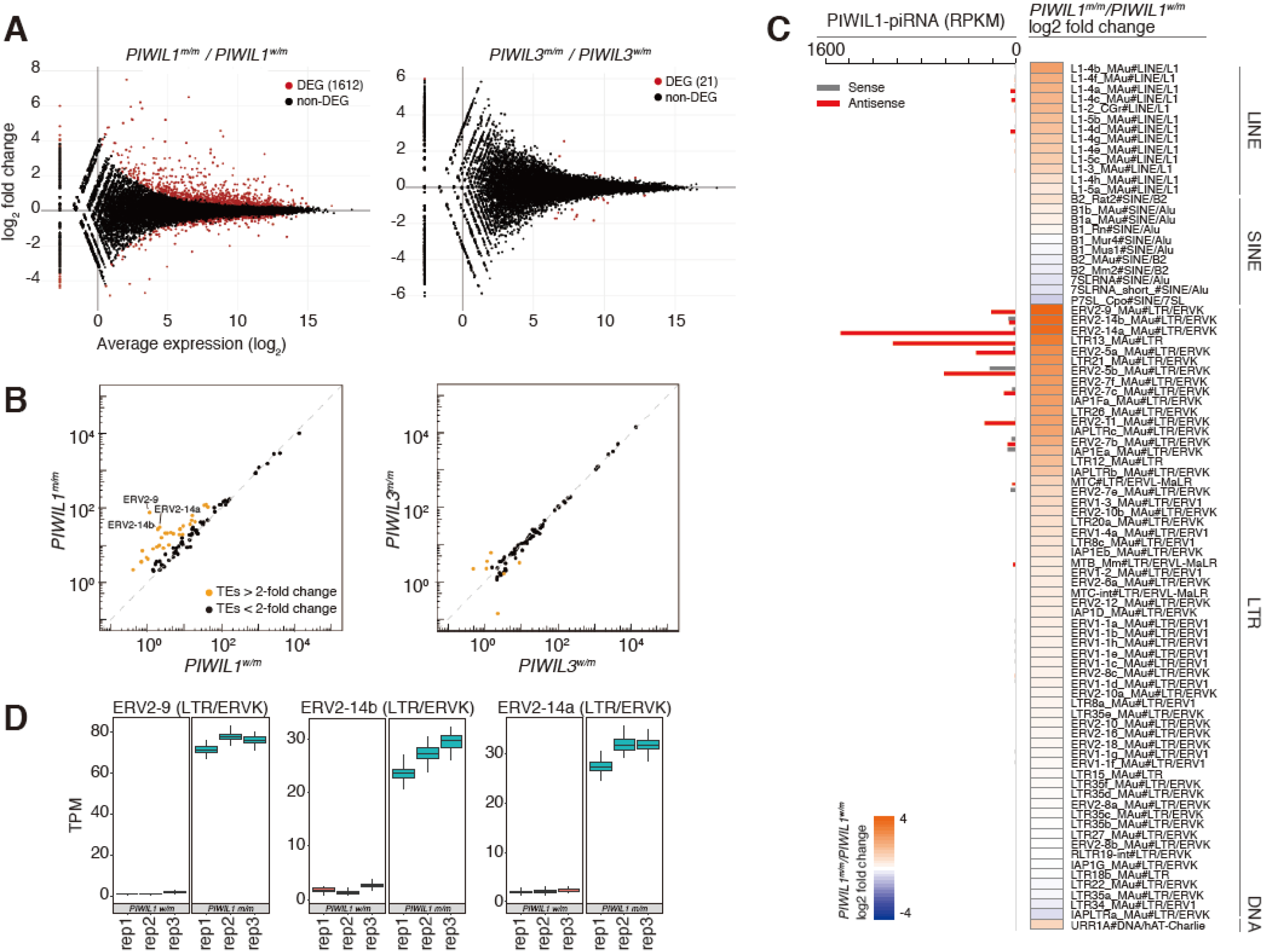
Silencing of a number of genes and TEs requires *PIWIL1*, but not *PIWIL3*, in golden hamster MII oocytes. **(A)** An MA plot showing the average expression level and log2 fold change of transcriptome in *PIWIL1* mutants (left) and *PIWIL3* mutants (right). Identified DEGs are indicated as red dots. **(B)** Dot plot showing the expression levels of TEs in *PIWIL*1 mutants (left) and *PIWIL3* mutants (right). Orange dots indicate TEs with changes in expression level >2-fold. **(C)** TE families with expression level >2 TPM in either *PIWIL1*^m/m^ or *PIWIL1*^w/m^ samples, and >0 TPM in both *PIWIL1*^m/m^ or *PIWIL1*^w/m^ samples, are listed with *PIWIL1*^m/m^/*PIWIL1*^w/m^ expression level log2 fold change and PIWIL1-piRNA levels. Increased expression levels are indicated in orange, whereas decreased expression levels are indicated in blue in the heatmap. piRNA levels are shown as a bar graph, with the indicated small RNAs mapped to either the sense (gray) or antisense (red) directions of TEs. **(D)** Examples of up-regulated TE families in *PIWIL1* mutant oocytes in three replicate samples.

The impact of the loss of *PIWIL1* or *PIWIL3* on the expression level of TEs was further analyzed using RNA-seq. We recently re-sequenced the golden hamster genome and detected species-specific TEs (18). Notably, ∼80% of the expressed TEs (TPM >1) were newly detected members of the TEs. Analysis of the TE expression levels revealed that the loss of *PIWIL1* resulted in the increased expression of 29 families of TEs (> 2-fold). In sharp contrast, the loss of *PIWIL3* had little impact on the expression levels of TEs (Fig. 3B and Fig. S6). In *PIWIL1*-deficient oocytes, TE family members, including ERV2-9, ERV2-14b, and ERV2-14a, were up-regulated the most (Fig. 3C and D). Of the 29 up-regulated TE families, 18 were LTR and 11 were LINE TEs. In addition, 27 out of the 29 up-regulated TE families were golden hamster specific TEs (18), suggesting that *PIWIL1* regulates active TEs that were recently added to the golden hamster genome. We then compared the abundance of the PIWIL1-piRNA population corresponding to each TE family with the change in expression of that TE in *PIWIL1*-deficient oocytes (Fig. 3C). This revealed a correlation between the abundance of PIWIL1-piRNAs antisense to particular TEs and up-regulation of these TEs with a lack of *PIWIL1*. Together, these results show that *PIWIL1* can regulate recently identified active TEs via their targeting by piRNAs in oocytes.

We performed gene ontology (GO) enrichment analysis of the DEGs identified with *PIWIL1* deficiency and found enrichment of terms related to nucleosome assembly and transcriptional and epigenetic regulation (Fig. S7A and B). This suggests that a loss of *PIWIL1* may cause defects in chromatin and/or the genome integrity network. Consistent with this notion, *PIWIL1*-deficient 2C nuclei displayed a single enlarged nucleolus with altered nuclear DNA enrichment, while 2C nuclei of *PIWIL1* heterozygous and *PIWIL3* homozygous oocytes displayed multiple nucleoli (Fig. S7C and D). This suggests that a loss of *PIWIL1* may induce the nucleolar stress response, which is often associated with cell cycle arrest and cell death (20, 21).

In mice, nuclear *Miwi2*/*Piwil4* acts as an effector for *de novo* DNA methylation of target TEs in embryonic male germ cells (7, 22, 23). Recent studies indicated that other PIWI proteins, which are predominantly cytoplasmic, may also function in DNA methylation in a *Miwi2*-independent manner in male germ cells (24, 25). These findings prompted us to examine DNA methylation in *PIWIL1*- and *PIWIL3*-deficient oocytes, although both PIWIL1 and PIWIL3 are predominantly cytoplasmic in oocytes (18) (also see Fig. S3B). Because it is known that de novo DNA methylation is complete by the germinal vesicle (GV) stage in mouse oocytes (26), we stained hamster *PIWI* mutant and control GV oocytes with an anti-5-methylcytosine (5mC) antibody. The DNA methylation level was approximately the same in *PIWIL1*-deficient and control GV oocytes (Fig. 4A and Fig. S8). However, *PIWIL3*-defficient oocytes had a significantly reduced DNA methylation level compared to control oocytes (Fig. 4B and Fig. S8). This suggests the involvement of *PIWIL3* in DNA methylation during oocyte development.

**Fig. 4.**
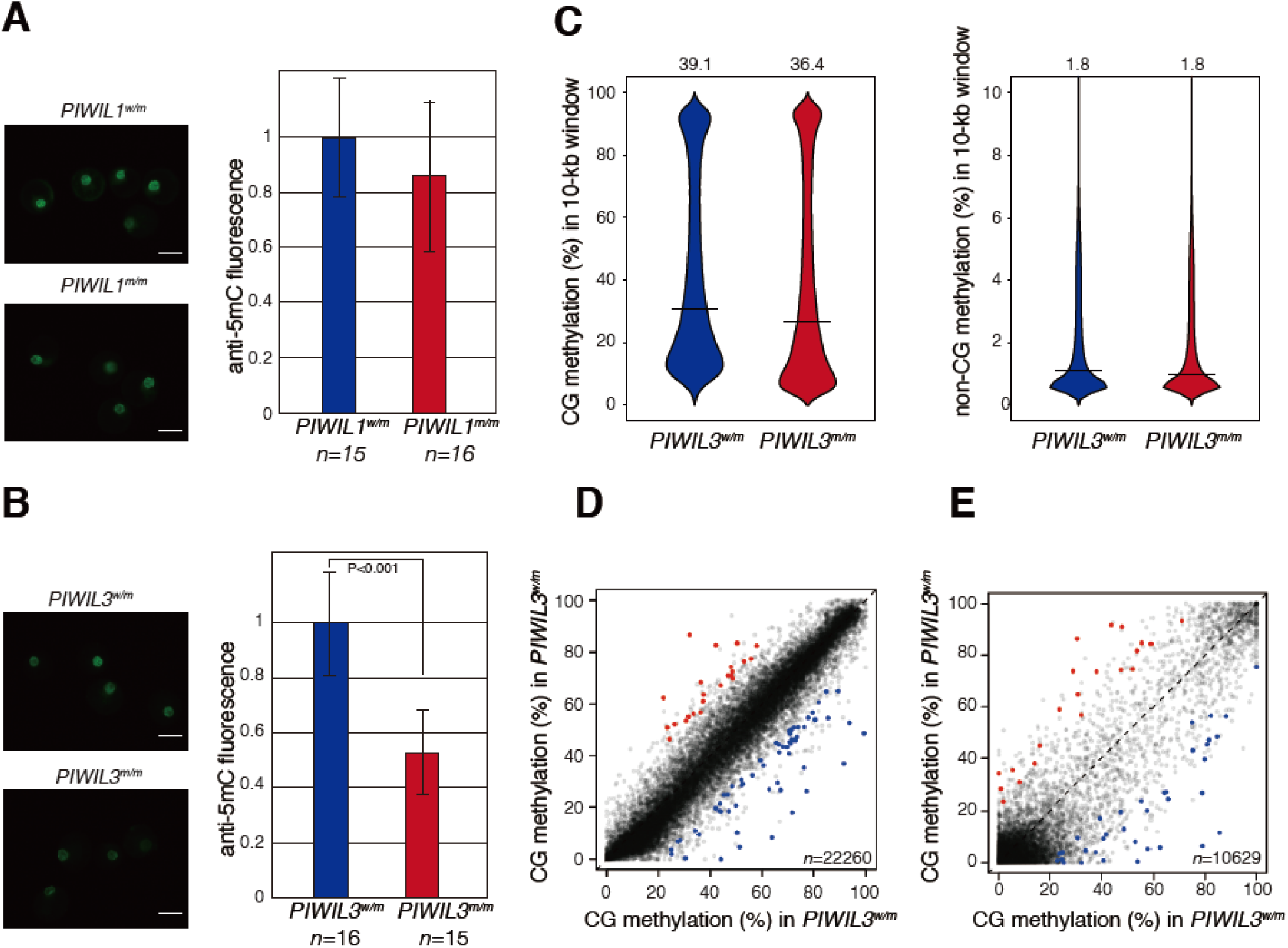
Loss of PIWIL3 leads to changes in DNA methylation in GV oocytes of golden hamsters. **(A, B)** 5mC staining of GV oocytes of *PIWIL1* **(A)** and *PIWIL3* mutants **(B)**. The amounts of 5mC were determined by measuring the fluorescence intensity using image J software (right). *n* shows the number of GV nuclei analyzed, and error bars indicate ± s.d. **(C)** A 10-kb window (*n*=230,285) violin plot of CG and non-CG methylation analyzed by whole genome bisulfite sequencing. Crossbars indicate the median value. The global CG and non-CG methylation levels were indicated above each graph. **(D, E)** Scatterplots describing the CG methylation levels in individual gene bodies **(D)** and gene promoters **(E)** in GV oocytes. Differentially methylated genes were defined as genes with >20% CG methylation differences between heterozygous and homozygous mutants, and p values between all heterozygous and homozygous replicates were <0.05. Among the differentially methylated genes, significantly less methylated genes were plotted in blue, whereas more methylated genes were plotted in red. p values were calculated by t test.

To explore the DNA methylation status in greater detail, we performed whole genome bisulfite sequencing. This analysis revealed that the level of global CG methylation (DNA methylation at CG sites, which are the major methylation target) was significantly decreased in *PIWIL3*-defficient GV oocytes compared to that in heterozygous control oocytes (36.4% versus 39.1%, respectively) (Fig. 4C). Our recent study showed that piRNA loading onto PIWIL3 is visible only in ovulated oocytes (18), suggesting a piRNA-independent role of PIWIL3 in DNA methylation. The level of non-CG methylation was unchanged. Measurement of CG methylation at piRNA clusters, TEs, piRNA target TEs and satellite repeats revealed that their total CG methylation level was not significantly altered in *PIWIL3* mutants (Fig. S9A–C).

Unlike any other differentiated mammalian cells, oocytes have DNA methylation predominantly within gene bodies with almost no methylation occurring in intergenic regions (24, 25). Therefore, we analyzed the DNA methylation status of gene bodies and gene promoters in *PIWIL3*-deficient oocytes and found that a number of genes showed altered levels of CG methylation at their gene bodies or promoters, with a larger population having reduced methylation, in *PIWIL3*-deficent GV oocytes (Fig. 4D and E). In addition, although there was no significant difference in oocyte transcripts between *PIWIL3* mutant and controls (Fig. 3A), genes having increased DNA methylation at their gene bodies tended to be expressed at higher levels (Fig. S9D). These results show that *PIWIL3* plays a role in DNA methylation in oocytes and suggest that the 3D chromatin structure could be altered in *PIWIL3*-deficient oocytes. The significant decrease in 5mC staining intensity in *PIWIL3* mutant GV oocytes (Fig. 4B and Fig. S8) may also be explained by possible changes in antibody accessibility due to altered chromatin structure. We found that meiosis occurs earlier in *PIWIL3*-deficient oocytes (Fig. S10A), suggesting either premature entry into meiosis or accelerated post-ovulatory aging. The morphology of the MII spindle was also altered in *PIWIL3*-deficient oocytes, with wider metaphase plates and longer acute-angled poles observed (Fig. S10B). Together, these findings suggest that the loss of *PIWIL3* may affect chromosome segregation in oocytes and early embryos, accounting for the appearance of three-cell or five-cell embryos that lack *PIWIL3* (Fig. 1D).

## Discussion

The establishment of DNA methylation in mouse oocytes and sperm is primarily due to the activity of DNMT3A and its cofactor DNMT3L (28–31). *DNMT3A*- and *DNMT3L*-deficient mouse oocytes can be fertilized normally but embryos derived from these oocytes die by E10.5, largely due to the absence of DNA methylation at maternally imprinted regions, leading to the notion that the only definitive roles for oocyte DNA methylation are in post-fertilization contexts (28–30). Nearly half of the embryos derived from *PIWIL3*-deficient hamster oocytes have a 2C block phenotype, but the observed DNA hypomethylation alone cannot account for the observed phenotype. Since transcripts from both TEs and protein-coding regions were not significantly affected in *PIWIL3*-deficent MII oocytes, *PIWIL3* may be involved in the formation of developmentally competent chromatin, which could be inherited by the zygote.

We analyzed *PIWIL1-* and *PIWIL3*-deficient hamsters and showed that maternally expressed *PIWIs* are important for the development of preimplantation embryos. *PIWIL1* regulated the expression of genes, including transposons, while *PIWIL3* had completely different functions, such as DNA methylation, meiosis and cell division. The origins of the piRNAs that bind to each PIWI are almost the same (18), suggesting that the length of the piRNA, the PIWI-piRNA complex, or PIWI protein itself was responsible for its respective biological characteristics.

Although mice have contributed immensely to our understanding of the physiology of humans, it is also increasingly clear that the genetic and physiological differences between humans and mice hamper the extrapolation of the results obtained in mouse models to direct applications in humans. We infer that the emergence of an oocyte specific Dicer isoform (32) could reduce the impact of the *Piwi* genes on regulation of gene expression, leading to the lack of the dependence on the piRNA pathway in the female germline of mice. The lack of discernible abnormalities in *Piw*i-deficient female mice may represent a special case of the piRNA pathway in mammals. Indeed, our study demonstrates that *PIWI*-deficient hamsters show abnormalities in oocyte function. This also suggests that *PIWI* genes should be studied as human infertility genes in women.

## Acknowledgments

We thank all members of the Siomi laboratory, especially H. Ishizu and K. Murano, for discussions and comments on this work, I. Ishimatsu for preparing paraffin sections, and P. Svoboda (IMG, Czech Academy of Sciences) and J. Li (Model Animal Research Center, Nanjing Medical University) for sharing unpublished data. We also thank J. Oishi of the Sasaki laboratory and T. Akinaga (Laboratory for Research Support, Medical Institute of Bioregulation, Kyushu University) for their help with whole-genome bisulfite sequencing. We are also grateful to M. Okabe and M. Ikawa (Osaka University) for their comments on the manuscript.

## Funding

Supported by Grant-in-Aid for the Japan Society for the Promotion of Science (JSPS) KAKENHI Grant Number 20H03175 (H.H), by Takeda Science Foundation (H.H), by JSPS Fellows (18J22025) (K.I), by funding from JSPS KAKENHI Grant Numbers 19H05268 and 18H02421, and from JST PRESTO Grant Number JPMJPR20E2 (Y.W.I), for Scientific Research (S) (<u>25221003</u>) and the Program of totipotency (<u>19H05753</u>) (H. Siomi). H. Siomi is also a recipient of funding for the project for elucidating and controlling mechanisms of aging and longevity (<u>1005442</u>) from the Japan Agency for Medical Research and Development (AMED). This work was also supported by JSPS KAKENHI Grant Number JP18H05214 (H. Sasaki).

## Author contributions

H. Siomi and H. H. designed the experiments. H. H. generated the *PIWIL1*- and *PIWIL3*-deficient hamsters with the help of H. M. K. I., and Y. W. I. conducted small RNA-seq and RNA-seq and analyzed the obtained data. A.Y. W. K., and H. Sasaki conducted whole-genome bisulfite-seq and analyzed the obtained data. H. Siomi, H. H., Y. W. I and H. Sasaki wrote the paper.

## Competing interests

The authors declare no competing interests.

## Data and materials availability

Raw sequence data generated during this study are available in the Gene Expression Omnibus (GEO) repository under accession no. GSE164356. Requests for materials should be addressed to H. Siomi. All data presented in this study are available in the main text or the supplementary materials.

## Notes

### Competing Interest Statement

The authors have declared no competing interest.

