## Supplementary Materials for "Production of functional oocytes requires maternally expressed *PIWI* genes and piRNAs in golden hamsters"

**This PDF file includes:**

Materials and Methods
Figs. S1 to S10
Tables S1

### Materials and Methods

#### Animals

Golden hamsters were purchased from Japan SLC and maintained under 14-h light/10-h dark cycles. All animal experiments were approved by the Animal Care and Use Committee of the Keio university.

#### Collection of GV oocytes, MII oocytes, PN zygotes and 2-cell embryos

Mature females were induced to superovulate by intraperitoneal (i.p.) injection of 40 IU PMSG (ASKA Animal Health) between 12:00 and 14:00 on the day of conspicuous, post-estrus vaginal discharge (day 1 of the estrous cycle). GV oocytes were collected from the ovary at 102h after PMSG injection. MII oocytes were collected from the oviduct at 115 h after PMSG injection. PN zygotes were collected from the oviduct at 91 or 115 h after PMSG injection followed by mating with males during the night of day 4 or 5 (depending on the female estrus cycle). Two cell embryos were collected from the oviduct at 121 or 145 h after PMSG injection, followed by mating with males during the night of day 4 or 5. Oocytes and embryos were collected from ovaries or oviducts with warmed and equilibrated HECM-10 medium (1) and covered with paraffin oil (Nakarai). All embryos were then washed twice to forth and cultured at 37.5°C under 5% $CO_2$. Culture dishes were pre-equilibrated for at least 5 h before use. The experiments were performed in a dark room with a >600 nm light.

### Preparation of sgRNA and Cas9

The CRISPR/Cas9 target sites of *PIWIL1* were designed using CRISPRdirect (http://crispr.dbcls.jp/) and produced using PCR-amplified templates and MEGA script T7 (Thermo Fisher Scientific). The *PIWIL3* sgRNAs were designed using CRISPRdirect (http://crispr.dbcls.jp/) and synthesized Alt-R CRISPR-Cas9 sgRNA (IDT) and annealed with Alt-R® CRISPR-Cas9 tracrRNA (IDT). Cas9 was injected either as mRNA or protein. Cas9 mRNA was transcribed from the T7 promoter-tagged Cas9 gene amplified from pX330 using the mMESSAGE mMACHINE™ T7 Transcription Kit and Poly(A) Tailing Kit (Thermo Fisher Scientific). Cas9 protein was purchased from IDT. All primers are listed in Table S1.

### Generation of *PIWIL1* and *PIWIL3* mutant golden hamsters

The technique for PN microinjection in golden hamsters is the same as that for mice, but with embryo manipulation conditions specifically developed for hamsters. In brief, fully equilibrated HECM-10 (37.5°C, 5% $CO_2$) covered with mineral oil was used as the injection medium. Embryos from one female were transferred to a 10 µL HECM-10 drop on the microinjection dish and 600 nM CRISPR-RNA and 200 nM Cas9mRNA (*PIWIL1*) or chemically synthesized 1 µM sgRNA and 1 µM Cas9 protein complex (*PIWIL3*) solution was injected into the pronucleus and/or cytoplasm of an embryo. Microinjections were performed on a heated microinjection stage (37°C) with red filters (cut off <600nm) and all embryo handling procedures were performed in a dark room. After injection, embryos were transferred to the oviducts of pseudopregnant females within 30 min. Recipients were allowed to naturally deliver and raise their pups.

**Establishment of *PIWIL1* and *PIWIL3* mutant lines**

Founder pups were mated with wild type males or females to produce the F1 generation. Genomic DNA was extracted from small pieces of tail tissue from F1 pups. Approximately 1-kb genomic fragments containing the target site were amplified by PCR using specific primers, and direct sequencing was performed. All primers are listed in Table S1. The *PIWIL1*-deficient mutant line ($15A^{del4}$) and *PIWIL3*-deficient mutant line ($26A^{del192}$) used in the present study were mated at least five times with wild types to avoid off-target effects. *PIWI* mutant golden hamster lines were maintained by mating with a wild type purchased from a breeder once every six months, because golden hamsters are a closed colony.

**Immunofluorescence**

The ovaries from wild-type and mutant female golden hamsters were fixed with 4% paraformaldehyde for preparing paraffin-embedded sections. Paraffin-embedded tissue blocks were cut into 3-µm slices, heated with antigen retrieval buffer (100x Tris-EDTA Buffer, pH 9.0, Abcam) in a microwave for 2 min after boiling and treated with anti-PIWIL1 monoclonal antibody (1A5) (2), or anti-PIWIL3 monoclonal antibody (3E12) (3). An Alexa488-conjugated goat anti-mouse IgG (Thermo Fisher) was used as the secondary antibody. For staining the acrosome, 3-µm paraffin-embedded testis sections were treated with PNA-FITC (Sigma-Aldrich). To visualize genomic DNA, the sections were counterstained with DAPI (DOJINDO). Fluorescence was observed using an IX71 fluorescence microscope (Olympus).

### Western blotting

Lysates of the MII oocytes were prepared by direct lysis with 2x SDS protein sample buffer. Total protein from 20 MII oocytes was separated on SDS-polyacrylamide gels and transferred to PVDF membranes. PIWIL1, PIWIL3 and TUBB were visualized using an ECL detection system (GE Healthcare) after incubation with anti-PIWIL1 monoclonal antibody (1A5), anti-PIWIL3 monoclonal antibody (3E12) or anti-TUBB monoclonal antibody (DSHB, E7). Horseradish peroxidase-linked sheep anti-mouse IgG (MP BIOMEDICALS) was used as the secondary antibody.

### Fertility analysis

Proestrus stage of 2- to 4-month-old mutant female golden hamsters were caged overnight with males at a 1:1 ratio, and vaginal sperm was examined the following morning. The pregnancy rate was calculated according to the number of successful deliveries per observation of sperm in the vagina.

### Small RNA sequencing

Total RNA was extracted from MII oocytes of *PIWIL1* and *PIWIL3* homozygous and heterozygous mutants for small RNA sequencing. The construction of small RNA libraries was performed using the NEXTFLEX® Small RNA-Seq Kit v3 (PerkinElmer) according to the manufacturer's instructions. The small RNA libraries were sequenced using the MiSeq (Illumina) platform, with three replicates from different samples sequenced for each mutant.

**Small RNA-seq data analysis**

Small RNA-seq analysis was performed as previously described (4) with slight modifications. Briefly, the adapter sequence and 4nt random sequences at the 3' and 5' ends were removed using cutadapt v1.14 (9). A quality filter of 30 was adopted by filtering sequences below this threshold. Reads in the range of 10–40 nt after adapter removal were used for further analysis. Reads were mapped to the hamster reference genome (hamster.sequel.draft-20200302.arrow.fasta) (3) using Bowtie (v1.1.1) (10), permitting 0 mismatches, and genome-mapped reads were used for further analysis. Genome-mapped reads were compared between replicate samples (n=3 per sample) and combined because of their high reproducibility ($r^2$ = 0.71 ~ 0.95). Reads were further mapped to known tRNA, rRNA, sn/snoRNA, and miRNA sequences. The tRNA sequences were obtained from RepeatMasker, while rRNA, sn/snoRNA, and miRNA sequences were obtained from Ensemble BioMart (11). The read count mapped to miRNAs was used to normalize small RNA reads obtained from heterozygous and homozygous mutants. Read counts mapped to each unique small RNA sequence were compared between heterozygous and homozygous samples, and small RNAs identical to the PIWIL1-bound piRNAs/PIWIL3-bound piRNAs (3) and those that decreased by over 4-fold in homozygous mutants were defined as PIWIL1-piRNAs or PIWIL3-piRNAs. The reads mapped to known ncRNAs were removed from the PIWIL1- and PIWIL3-piRNAs. Nucleotide composition was analyzed using WebLogo v3.7.5 (12), and annotation of the PIWIL1- and PIWIL3-piRNAs were assigned using a previously described script (5). The priority of the assignment was in the order of transposons, repeats, and protein-coding genes. The reads were assigned to the highest priority annotation, and therefore every read was assigned to a single annotation. PIWIL1- and PIWIL3-piRNAs were further mapped to the consensus sequence of transposon families obtained from the RepeatMasker database and our previous research (3). The mapping to transposon consensus sequence was performed using Bowtie (v1.1.1) (10), with up to two mismatches permitted. Mapped reads were counted and normalized to the total number of mapped reads and the length of each transposon sequence.

**RNA-seq of MII oocytes**

Total RNA was purified from 50 MII oocytes of 2- to 4-month-old golden hamsters using the NucleoSpin RNA Plus XS kit (MACHEREY-NAGEL) for RNA-seq. RNA-seq libraries were constructed using total RNA from MII oocytes and the SMART-Seq Stranded Kit (TaKaRa Bio). Sequencing was performed using the NovaSeq 6000 (Illumina) platforms.

**RNA-seq data analysis**

Prepared libraries were sequenced in paired ends, and triplicate samples were obtained per experiment. Low-quality sequences (quality value lower than 30) and 3-nt random sequences were removed using cutadapt v1.14 (9) software. Mapping to the golden hamster genome (hamster.sequel.draft-20200302.arrow.fasta) (3) was performed using HISAT2 (13) with default parameters. featureCount v2.01 (14) was used to count reads mapped to genes. featureCounts output was further processed using the TCC-GUI version 2019.02.08 (15) with default parameters. The TCC-GUI uses the TMM normalization method, and edgeR was used to filter differentially expressed genes (DEGs). Extracted DEGs were converted to mouse homolog genes using Ensemble BioMart (11), and DAVID (16) was used to perform gene ontology (GO) enrichment analysis. Salmon v1.3.0 (17) and Sleuth v.0.30.0 (18) were used for quantification of the expression levels of TEs. Wasabi v.1.0.1 was used to analyze the Salmon output in Sleuth. To map the TE reads, consensus sequences of TEs were obtained from RepeatMasker and our previous analysis (3), and the TE sequences were merged with the other gene sequences. The mean value for three replicates was used to determine the overall expression levels of TEs.

**MII spindle staining**

MII oocytes were freed from the zona pellucida by treatment with acidic Tyrode's solution. Zona-free GV oocytes were fixed using 4% paraformaldehyde for 10 min and permeabilized with 0.1% Triton X-100 in 0.01% PVA/PBS for 2 min at room temperature, respectively. For immunofluorescence staining, we used Alexa 488 conjugated tubulin alpha monoclonal antibody (MBL). Cells were incubated at 4°C overnight with a primary antibody and then washed three times. To visualize genomic DNA, the cells were counterstained with DAPI (DOJINDO). Fluorescence was measured using an IX71 fluorescence microscope and FV3000 laser scanning confocal microscope (Olympus).

**5mC staining**

GV oocytes were freed from the zona pellucida by treatment with acidic Tyrode's solution.
Zona-free GV oocytes were fixed using 4% paraformaldehyde for 10 min and permeabilized with 0.1% Triton X-100 in 0.01% PVA/PBS for 2 min at room temperature, respectively. These cells were treated with 2N HCl 10 min at 37°C and washed twice with 0.01% PVA/PBS. For immunofluorescence staining, we used a monoclonal primary antibody against 5mC, 33D3 (Abcam). Cells were incubated at 4°C overnight with the primary antibody and then washed and incubated with an Alexa Fluor 488-onjugated anti-mouse IgG secondary antibody (Thermo Fisher Scientific) at room temperature for 1 h. Fluorescence was measured using an IX71 fluorescence microscope (Olympus).

**Whole genome bisulfite sequencing and data analysis**

DNA from 30 GV oocytes was spiked with 1% unmethylated lambda phage DNA. Libraries were generated by the post-bisulfite adaptor tagging method with eight cycles of library amplification (6, 7) and sequenced using the NovaSeq 6000 platform (NVCS v1.6 and RTA v3.4.4) (8). Data analysis was performed as previously described (7). Briefly, reads were trimmed to remove low-quality bases, and were mapped to the golden hamster genome using Bismark v0.20.0. Methylation data at CG and non-CG sites covered with 10–500 reads were extracted and compared between the biological triplicates in 10-kb windows. After confirmation of the reproducibility (Pearson correlation coefficient >0.88 for *Piwil3*$^{w/m}$ and >0.92 for *Piwil3*$^{m/m}$), the triplicate data were combined and used for downstream analyses.

**A**

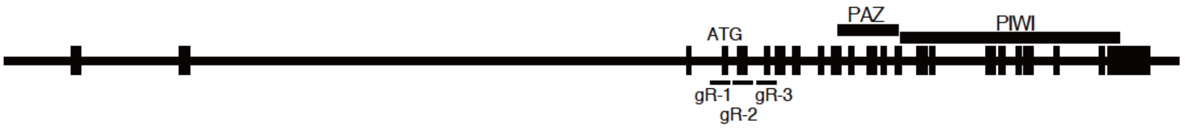

**B**

*PIWIL1* gR-1: AGGAGACGGTACAGCATGTT

*PIWIL1* gR-2: GACAGAGAGGAATGGTAGTT

*PIWIL1* gR-3: AGCTGTCATTGGCAGAGAGA

**C**

| | method | number of founder pups | number of mutant pups | mutant line |
|---|---|---|---|---|
| gR-1 | guide RNA + Cas9 mRNA | 16 | 7 | 15A$^{del4}$ |
| | guide RNA + Cas9 protein | 14 | 6 | 57A$^{del3}$ |
| gR-2 | guide RNA + Cas9 protein | 14 | 10 | 44A$^{del11}$, 44A$^{del15}$ 63A$^{del27}$, 63A$^{del3}$ 70A$^{del11}$, 70A$^{del15}$ |
| gR-3 | guide RNA + Cas9 protein | 2 | 0 | |

**D**

exon4

…tggcagAGG**ATG**ACGGGCCGAGCCCGAGCCCGGGCCCGCGGCAGGGCCCGCGGTC**AGGAGACGGTACAGCATGTT**GGGGCTGCTGCGgtgagt…

M T G R A R A R A R G R A R G Q E T V Q H V G A A A

exon5

…aagcagAGCCAGCAGCCTGGGTGCATCCCACCGAGACCTCAGCAGTCACCCGCAGAGGGGGACTTGGTTGGCCGAGGACGACAGAGAGGAATGGTAG…

S Q Q P G C I P P R P Q Q S P A E G D L V G R G R Q R G M V

**E**

exon4

…tggcagAGG**ATG**ACGGGCCGAGCCCGAGCCCGGGCCCGCGGCAGGGCCCGCGGTC**AGGAGACGGTACAGTT**GGGGCTGCTGCGgtgagt…

M T G R A R A R A R G R A R G Q E T V Q L G L L R

exon5

…aagcagAGCCAGCAGCCTGGGTGCATCCCACCGAGACCTCAGCAGTCACCCGCAGAGGGGGACTTGGTTGGCCGAGGACGACAGAGAGGAATGGTAG…

A S S L G A S H R D L S S H P Q R G T W L A E D D R E E W *

**Fig. S1 Strategy for the production of *PIWIL1* mutant golden hamsters by CRISPR/Cas9** **(A)** Structure of the golden hamster *PIWIL1* gene and target locus for sgRNA. **(B)** The guide RNA sequence for the golden hamster *PIWIL1* gene. **(C)** Production of *PIWIL1* mutant golden hamster lines. **(D)** The *PIWIL1* nucleic acid and amino acid sequences of the targeted site in wild-type hamsters. Upper case underlined denotes the exon sequence, bold with a red box denotes the target sequence, and red denotes the deleted sequence in the 15A$^{del4}$ line. **(E)** The *PIWIL1* nucleic acid and amino acid sequence of the targeted site in the mutant line (15A $^{del4}$). The amino acid sequences in red indicate sequences changed by deletion.

**A**

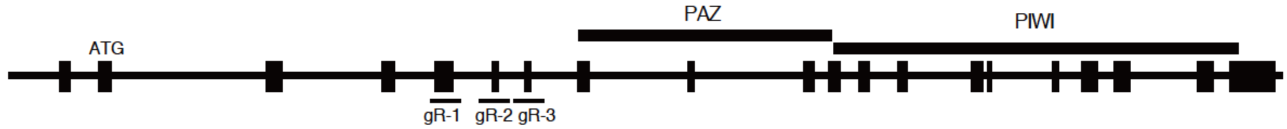

**B**

*PIWIL3* gR-1: ATGAGTTTCCATCGAAAATA

*PIWIL3* gR-2: TGTAATAGCGAAGCCACTCT

*PIWIL3* gR-3: AAGATGAGCTTATGGCGTCA

**C**

| | method | number of founder pups | number of mutant pups | mutant lines |
|---|---|---|---|---|
| gR-1 | guide RNA + Cas9 protein | 10 | 4 | |
| gR-2 | guide RNA + Cas9 protein | 4 | 4 | 25A$^{del89}$, 25B$^{del4}$ 25C$^{del300}$, 26A$^{del192}$ 26B$^{del41}$, 26C$^{del2}$ 26D$^{del39}$, 26E$^{in4}$ |
| gR-3 | guide RNA + Cas9 protein | 4 | 3 | 33A$^{del8}$, 33B$^{del172}$ 33C$^{del329}$, 36A$^{del8}$ 36B$^{del4}$ |

**D**

exon6

```
…tgaaatccttagGAAATGGAGCTTGTCAGTAAGACTGAAGGTGATGACAAAGTGAGAATCAGGATCCAGCTTTCCAGTGAACTCCACCC
             E  M  E  L  V  S  K  T  E  G  D  D  K  V  R  I  R  I  Q  L  S  S  E  L  H  P
CAGTCACCCAGAGTGGCTTCGCTATTACAACATCCTCTTTAGAAGgtgcatgagactttgtcttccccacccaccatgatccctggggtg
  S  H  P  E  W  L  R  Y  Y  N  I  L  F  R  R
ctcccccaccatggacgtgaacacagccacattccttcaaccgtgtgtgctctcaaggcctcatgttgttgagcagacaaggtcttctg
ggtaaaatgatatctgaacctcagctacaatgtgcaagttcggtatctaggccccagcatttgtcctcctgcctatacccacttcctgt…
```

**E**

exon6

```
…tgaaatccttagGAAATGGAGCTTGTCAGTAAGACTGAAGGTGATGACAAAGTGAGAATCAGGATCCAGCTTTCCAGTGAACTCCACCC
             E  M  E  L  V  S  K  T  E  G  D  D  K  V  R  I  R  I  Q  L  S  S  E  L  H  P
CAGTCACCtcagctacaatgtgcaagttcggtatctaggccccagcatttgtcctcctgcctatacccacttcctgt…
  S  H  L  S  Y  N  V  Q  V  R  Y  L  G  P  S  I  C  P  P  A  Y  T  H  F  L…
```

**Fig. S2 Strategy for the production of *PIWIL3* mutant golden hamster by CRISPR/Cas9**
**(A)** Structure of the golden hamster *PIWIL3* gene and target loci for sgRNAs. **(B)** The guide RNA sequences for the golden hamster *PIWIL3* gene. **(C)** Production of *PIWIL3*mutant golden hamster lines. **(D)** The *PIWIL3* nucleic acid and amino acid sequence of targeted site in wild-type hamsters. Upper case underlined denotes the exon sequences, bold with a red box denotes the target sequences, and red denotes the deleted sequence in the 26A$^{del192}$ line. **(E)** The *PIWIL1* nucleic acid and amino acid sequence of targeted site in the mutant line (26A $^{del192}$). The amino acid sequences in red indicate sequences changed by deletion.

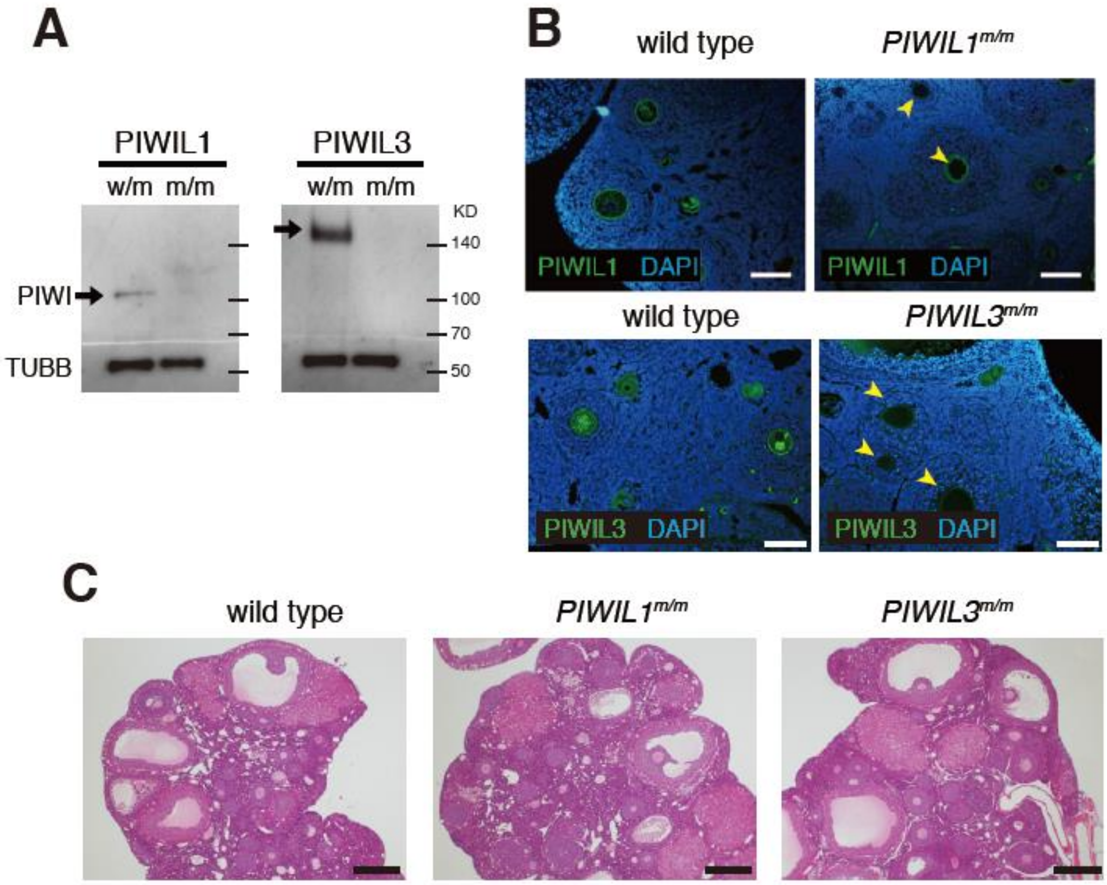

**Fig. S3 Lack of PIWI proteins in oocytes of *PIWIL1* or *PIWIL3* mutant hamsters**
**(A)** Total lysates of 20 MII oocytes each from heterozygous mutant (w/m) and homozygous mutant (m/m) golden hamsters were subjected to western blotting. TUBB was used as the loading control. Spectra™ Multicolor Broad Range Protein Ladder (Thermo Fisher Scientific) was used as size marker. **(B)** Immunological staining of *PIWIL1* and *PIWIL3* mutant ovaries with an anti-PIWIL1 and PIWIL3 antibodies. The arrowheads in the mutants indicate the ovarian follicles. The zona pellucida fixed with paraformaldehyde showed autofluorescence. In particular, PIWIL1, which requires a long exposure time, has strong ring-shaped fluorescence both in the wild type and *PIWIL1*$^{m/m}$. Scale bars = 50µm. **(C)** Ovarian sections from 8-week-old wild-type and *PIWIL1* and *PIWIL3* mutant golden hamsters. Hematoxylin and eosin-stained ovarian sections of wild-type, *PIWIL1* mutant and *PIWIL3* mutant golden hamsters (scale bars = 200 µm).

Note: By fixing PIWIs, the antigenicity is weakened. PIWIL1 antigenicity is particularly weak even after antigen retrieval treatment: therefore, the exposure time is long and the background of the zona pellucida becomes visible.

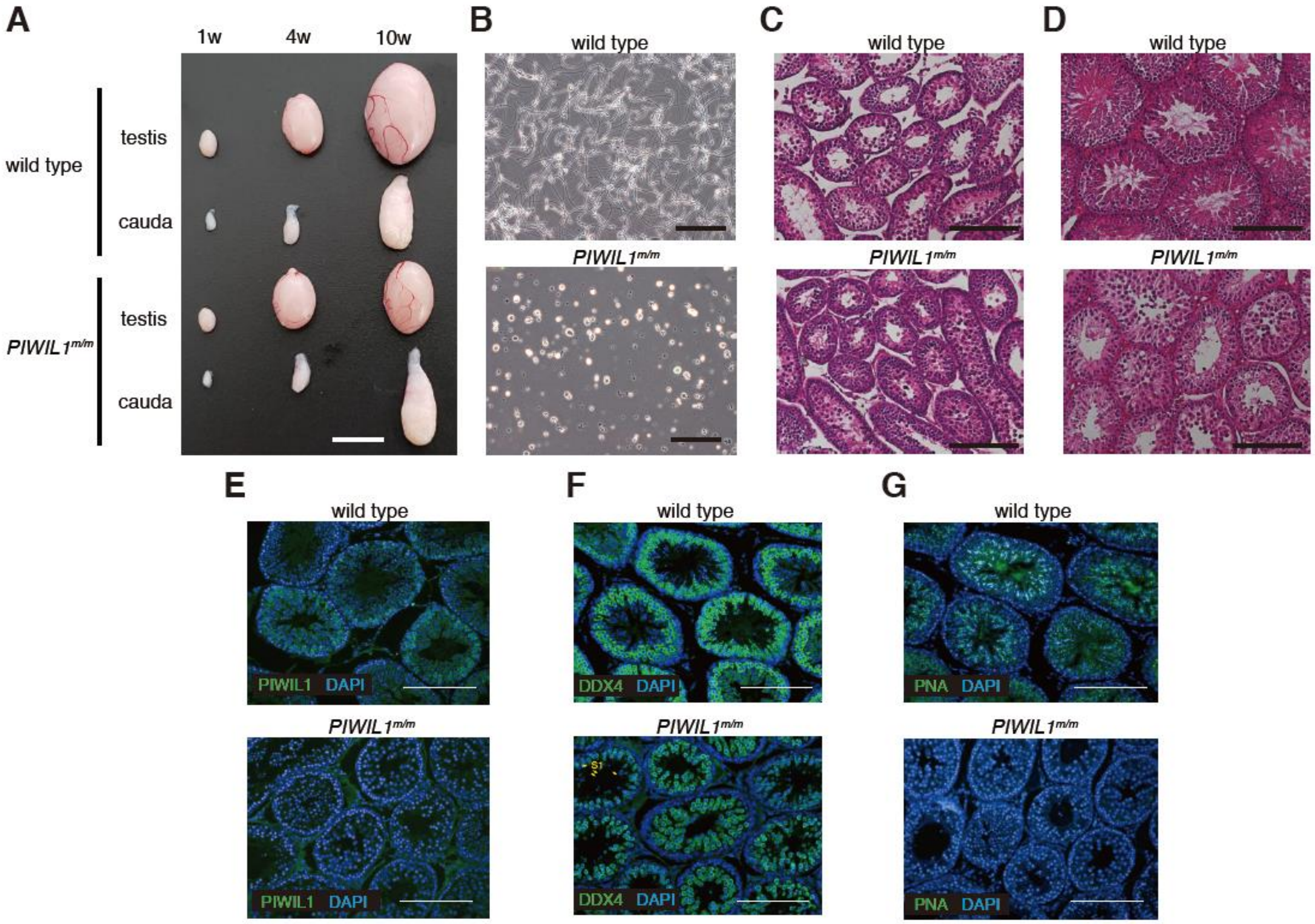

**Fig. S4 Defects in spermatogenesis in *PIWIL1* mutant male hamsters**

**(A)** Testis and cauda epididymis of 1week-old (1 w), 4-week-old (4 w) and 10-week-old (10 w) from wild-type and *PIWIL1* mutant golden hamsters (scale bars = 10 mm). **(B)** Sperm from the 10-w wild-type and *PIWIL1* mutant cauda epididymis. Scale bars = 50 µm. **(C, D)** Hematoxylin and eosin-stained 4-w **(C)** and 15-w **(D)** testis sections of wild-type and *PIWIL1*$^{m/m}$ hamsters, respectively. Scale bars = 100 µm. **(E, F, G)**; anti-PIWIL1 antibody **(E)**, anti-DDX4antibody **(F)** and PNA **(G)** staining of testis sections of 15-w wild-type and *PIWIL1* mutant hamsters. Almost no spermatids were observed after meiosis in the *PIWIL1* mutant. Scale bars = 100 µm.

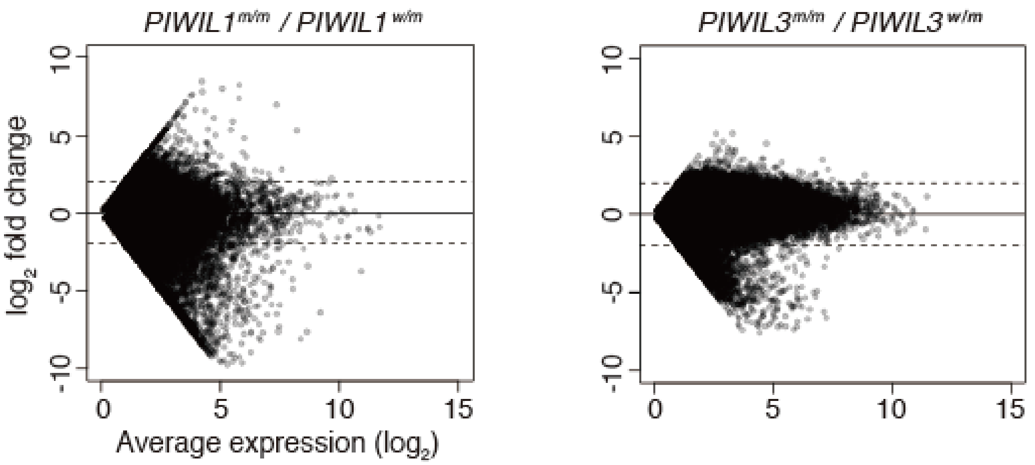

**Fig. S5 Changes in the expression levels of each unique small RNA sequence in *PIWI* mutants.**
An MA plot showing the expression level of each unique small RNA sequence. To calculate log2 fold change and expression levels, 1 was added to each value to calculate the levels of small RNA sequences with 0 in either sample.

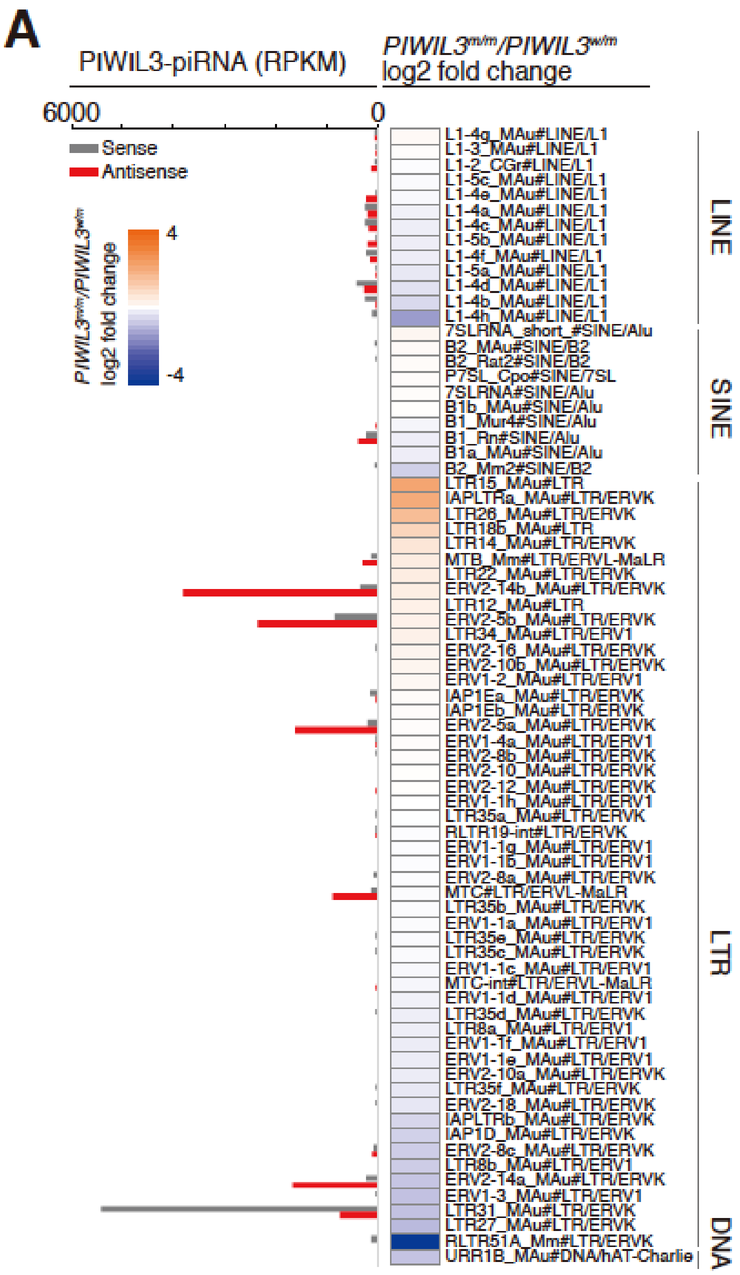

**Fig. S6 PIWIL3 and PIWIL3-piRNAs do not have a significant impact on TE silencing in MII oocytes**

TE families with expression levels >2 TPM in either *PIWIL3*$^{m/m}$ or *PIWIL3*$^{w/m}$ samples and >0 TPM in both *PIWIL3*$^{m/m}$ and *PIWIL3*$^{w/m}$ samples are listed with log2 fold change in *PIWIL3*$^{m/m}$/*PIWIL3*$^{w/m}$ expression level and PIWIL3-piRNA levels. Increased expression levels are indicated in orange, whereas decreased expression levels are indicated in blue in the heatmap. piRNA levels are shown as a bar graph, with indicated small RNAs mapped to either the sense (gray) or antisense (red) direction of TEs. PIWIL3 does not largely impact the expression levels of TEs, and TEs showing increased levels are not targeted by PIWIL3-piRNAs.

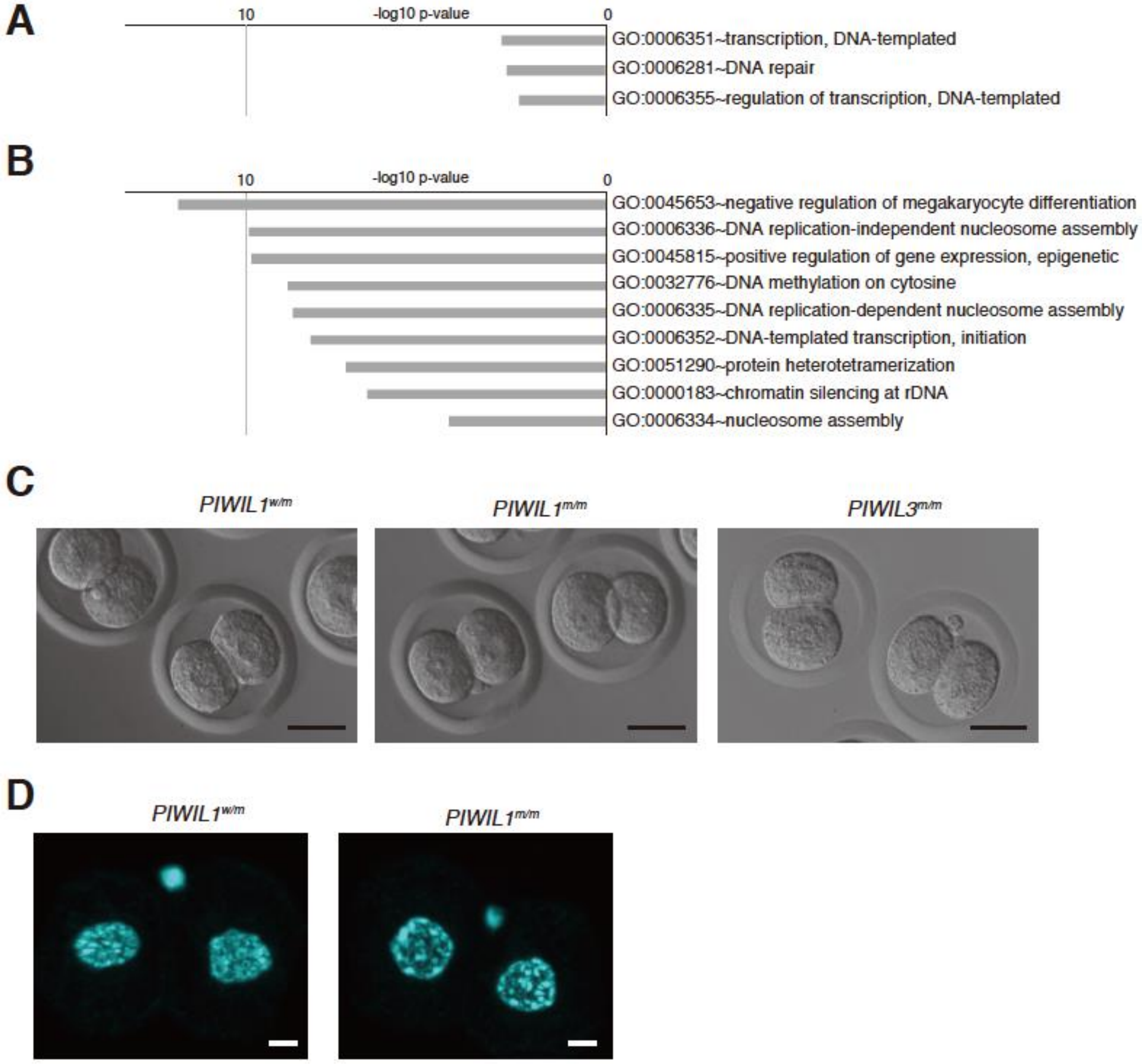

**Fig. S7 Gene ontology analysis and developmental abnormalities in *PIWIL1* mutant hamsters** **(A, B)** Enriched gene ontology (biological process) terms using DEGs obtained in *PIWIL1* mutant MII oocyte transcriptome analysis. **(A)** Upregulated genes in the *PIWIL1* mutants. **(B)** Downregulated genes in the *PIWIL1* mutants. Terms with p-values lower than 0.001 are listed. **(C)** Nucleus morphology of the 2C stage embryos. *PIWIL1*$^{m/m}$ showed a single large nucleolus, while *PIWIL1*$^{w/m}$ and *PIWIL3*$^{m/m}$ formed multiple small nucleoli. Scale bars = 50 µm. **(D)** The morphology of genomic DNA in 2-cell embryos was analyzed by DAPI staining. Scale bars = 10 µm.

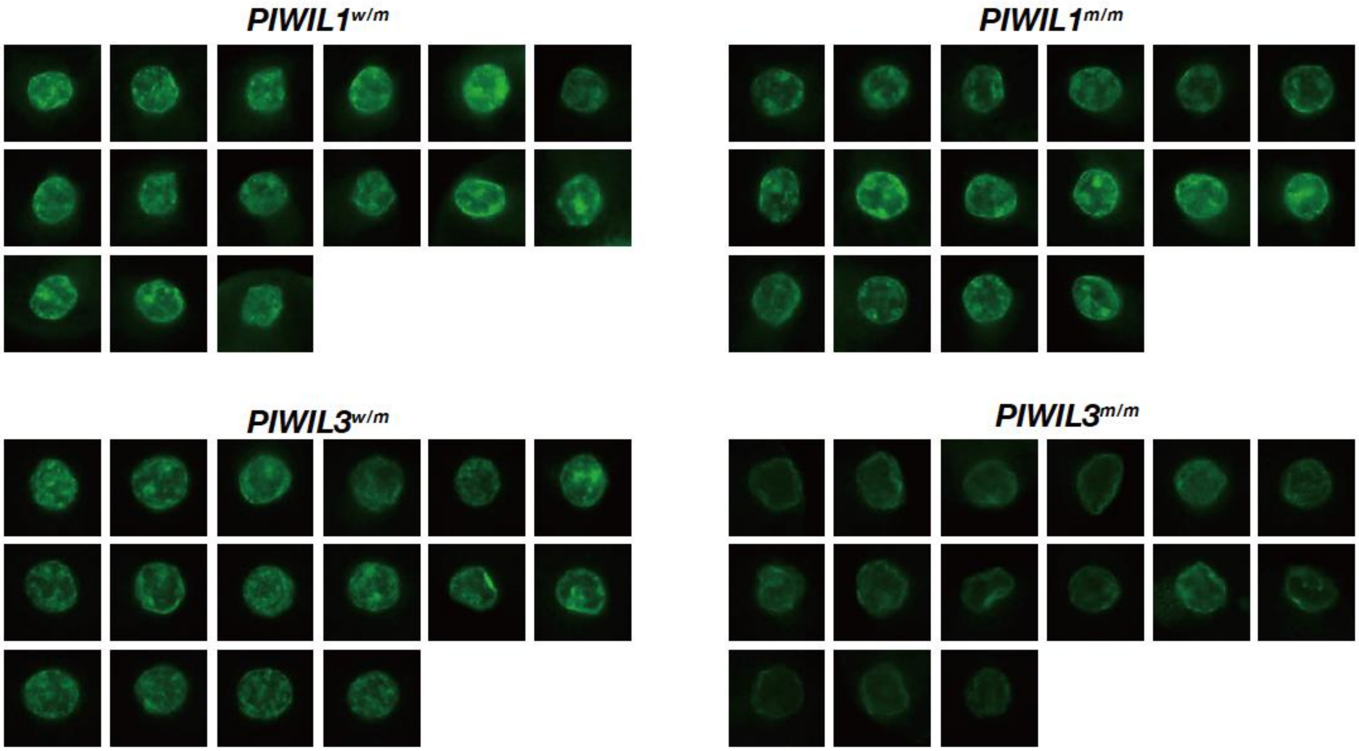

**Fig. S8 Representative GV oocytes stained with anti-5mC antibody**

GV oocytes were stained with a 5mC antibody (33D3) and the fluorescence GV nuclei were imaged. The fluorescence was analyzed by ImageJ and used to be described in Fig. 4A and B.

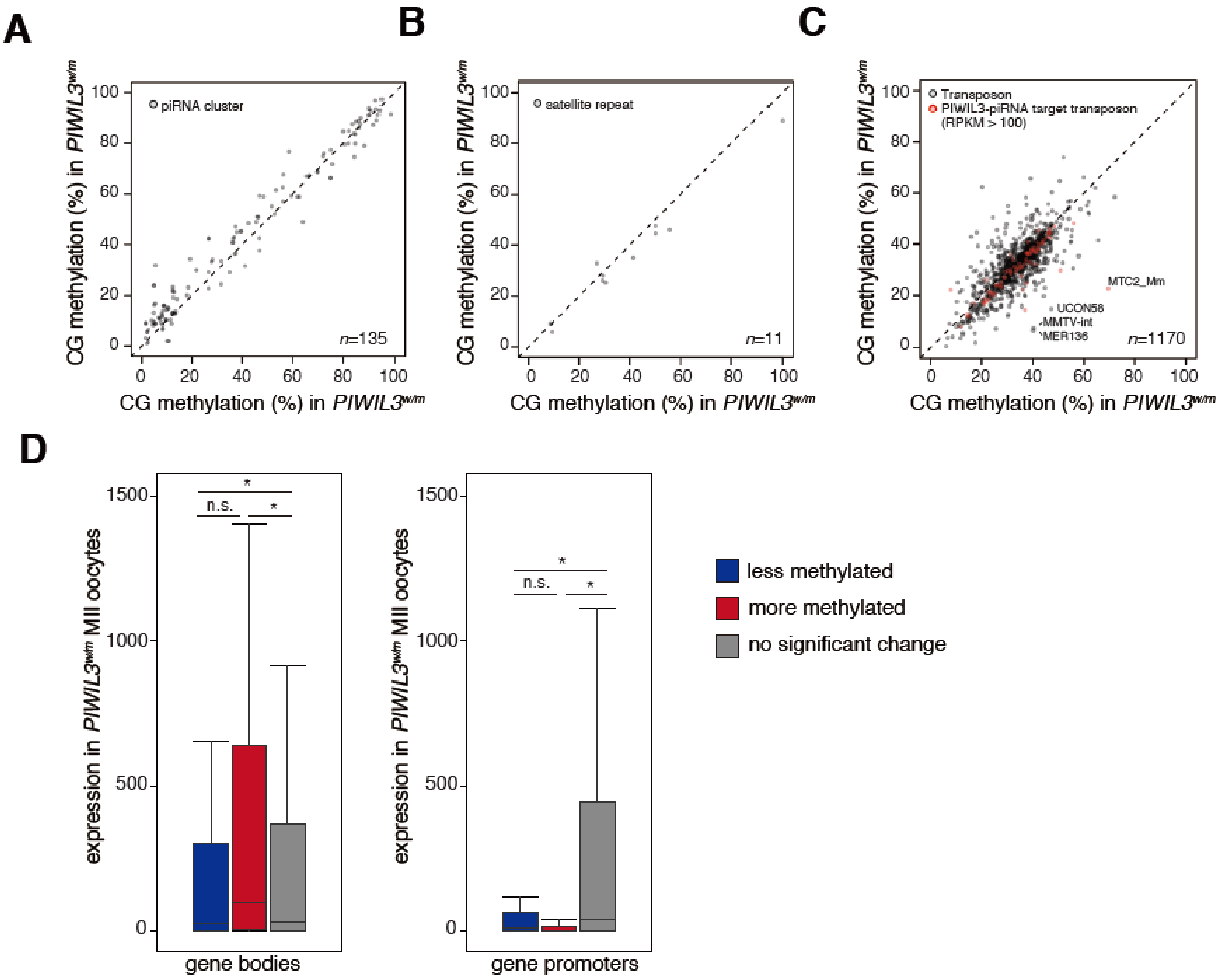

**Fig. S9 CG methylation of repeats in *PIWIL3* mutant GV oocytes**

**(A, B, C)** Scatterplot describing the CG DNA methylation levels of individual piRNA clusters **(A)**, satellite repeats **(B)**, and TE families **(C)** in GV oocytes. PIWIL3-piRNA target TEs (TEs with over 100 RPKM PIWIL3-piRNAs mapped) are plotted in red while the other transposons are plotted in gray (left). Names of differentially methylated TE families are indicated within the plots. **(D)** Boxplots showing the expression levels of genes with differential CG methylation found in their gene bodies or promoters. The left panel shows the expression levels of genes with altered methylation in gene bodies. The right panel shows the expression levels of genes with altered methylation in promoters. p values were calculated using the Mann Whitney U test.

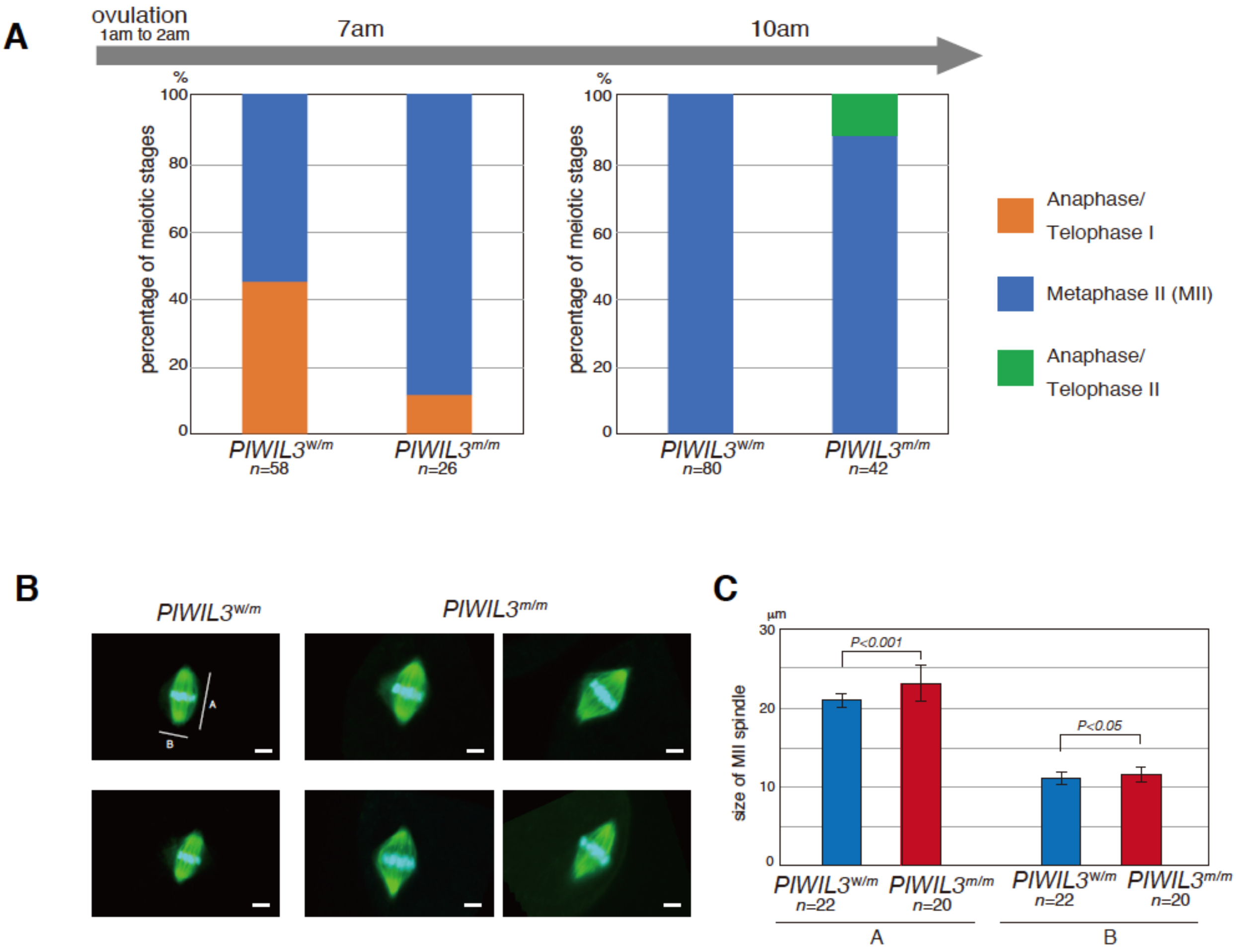

**Fig. S10 Meiosis and MII spindle formation in ovulated oocytes of *PIWIL3* mutant golden hamsters**

**(A)** *PIWIL3$^{w/m}$ and PIWIL3$^{m/m}$* oocytes were collected, and the stages of meiosis at 7 and 10 am in the morning were analyzed following TUBA and DAPI staining. **(B)** MII oocytes at 10 am in the morning were stained with anti-TUBA antibody (green) and DAPI (blue). Scale bars = 5 µm. **(C)** The size of the MII spindle (A and B of Fig. S10B) was measured and compared between *PIWL3$^{w/m}$ and PIWL3$^{m/m}$*. Error bars indicate ± s.d.

| | | |
|---|---|---|
| **Table S1. Pimers** | | |
| **for *PIWIL1* sgRNA and Cas9 mRNA synthesis** | | |
| | T7gR1_FW | GCGGCCTCTAATACGACTCACTATAGGGAGGAGACGGTACAGCATGTTGTTTTAGAGCTAGAAATAGCA |
| | gR1_RV | GCACCGACTCGGTGCCACTTTTTC |
| | T7mCas9_FW | GCGGCCTCTAATACGACTCACTATAGGGACCGGTGCCACCATGGACTAT |
| | mCas9_RV | CAGATGGCTGGCAACTAGAAGGC |
| **for genotyping of *PIWI1* mutant hamsters** | | |
| | PIWIL1#15 FW-M | GCGGTCAGGAGACGGTACAGTTGG |
| | PIWIL1#15 FW-W | GCGGTCAGGAGACGGTACAGCATG |
| | PIWIL1 G8 | CTGAGAGCTGGCAGTTGCCTGTGC |
| **for genotyping of *PIWI3* mutant hamsters** | | |
| | PIWIL3-6 | TGCTGACTGTTATTCAGTGGGACC |
| | PIWIL3-7 | AGCAGAAGCCAGAGATCTTGAGCC |
